# EEG-based speaker-listener neural coupling reflects speech-selective attentional mechanisms beyond the speech stimulus

**DOI:** 10.1101/2022.10.02.510499

**Authors:** Jiawei Li, Bo Hong, Guido Nolte, Andreas K. Engel, Dan Zhang

## Abstract

When we pay attention to someone, do we focus only on the sound they make, the word they use, or do we form a mental space shared with the speaker we want to pay attention to? Some would argue that the human language is no other than a simple signal, but others claim that human beings understand each other not only by relying on the words that have been said but also formed a shared ground in the specific conversation. This debate was raised early, but the conclusion remains vague. Our study aimed to investigate how attention modulates the neural coupling between the speaker and the listener in a cocktail party paradigm. The temporal response function (TRF) method was employed to reveal how the listener was coupled to the speaker at the neural level. The results showed that the neural coupling between the listener and the attended speaker peaked 5 seconds before speech onset at the delta band over the left frontal region, and was correlated with speech comprehension performance. In contrast, the attentional processing of speech acoustics and semantics occurred primarily at a later stage after speech onset and was not significantly correlated with comprehension performance. These findings suggest that our human brain might have adopted a predictive mechanism to achieve speaker-listener neural coupling for successful speech comprehension.

**Three key points:**

1. Listener’s EEG signals coupled to the speaker’s 5 s before the speech onset, which revealed a “beyond the stimulus” attentional modulation.
2. Speaker-listener attentional coupling is correlated to the listener’s comprehension performance, but the speech-listener’s coupling didn’t.
3. The implementation of temporal response function methods and the neural language methods yielded novel perspectives to the analysis of the inter-brain studies.

## Introduction

> *“Men do not understand one another by actually exchanging signs for things…they do it by striking the same note on their mental instruments.”*
>
> *Wilhelm von Humboldt*

When we pay attention to someone in a cocktail party, what sort of information from the speaker do the listeners exactly attend to? Are the listeners just perceiving sound waves or process semantic meaning in the speech that the speaker generated? Or do listeners and the speakers reach some common ground that is beyond the speech stimulus itself? The question is about the core connection formed between the speaker and the listener via speech. There are different perspectives on this issue: Some argued the speech is just a type of signaling (Chomsky, 1959; Skinner, 1957); others suggested that speakers and listeners need to be intentionally aligned, rather than relying solely on speech (Gadamer, 1975; Searle, 1980). The discourse has been primarily conducted at the theoretical level. However, empirical evidence, especially in neuroscience is still lacking to settle the debate.

The cocktail party paradigm together with the emerging inter-brain approach provides a unique and advantageous tool for examining this issue. The cocktail party established an ideal benchmark for contrasting the information that the listener deliberately focused on versus that which they disregarded (Cherry, 1953; McDermott, 2009; Middlebrooks *et al*., 2017; Shinn-Cunningham, 2008). In this situation, listeners would receive the information from different speakers simultaneously, but the listener only processes the speech that she or he attended in a cocktail party. The inter-brain method entails the acquisition of neural data from both the speaker and the listener and introduces the speaker’s neural activity as a new reference in the speech comprehension (Hasson *et al*., 2012; Jing Jiang *et al*., 2021; Z. Li *et al*., 2021; Pérez & Davis, 2023; Redcay & Schilbach, 2019; Schoot *et al*., 2016; Stolk *et al*., 2016). Using this method, the neural coupling with the speaker and the neural coupling with the speech could be compared simultaneously. If the listener only perceives the information she or he received in the speech as the behaviorism scientists suggested (Chomsky, 1959; Skinner, 1957, 1986), the speaker-listener coupling would be largely bounded with the neural coupling with the speech stimuli itself. In other words, the neural coupling with the speaker would occur at the similar time window or the similar frequency band with the neural coupling with the speech stimuli. However, if the listener coupled with the speaker “beyond the stimulus”(Hartley & Poeppel, 2020) or “extra-linguistic”(Schoot *et al*., 2016), the time and frequency bands for the attention modulation of the speaker’s neural activity may reveal a different pattern comparing the speech-listener’s coupling.

Previous inter-brain studies have provided potential evidence of the beyond the speech attention. In the monologue situation, inter-brain studies revealed the distinct pattern of the speaker-listener coupling comparing with the speech-listener coupling: the speaker-coupling coupling occurs before the speech onset (Dai *et al*., 2018; Jing Jiang *et al*., 2015; L. Liu *et al*., 2020; Stephens *et al*., 2010; Zheng *et al*., 2018) or on the different frequency bands (Pérez *et al*., 2017). However, studies on speaker-listener coupling in a cocktail party scenario remains sparse. There are few studies investigating the attentional speaker-listener coupling or entrainment (Dai *et al*., 2018; Kuhlen *et al*., 2012). Direct comparison of neural mechanisms underlying attention modulation in speaker-listener coupling and listener-stimuli coupling is still insufficient. Previous studies only compared the coupling between the speaker-listener and speaker-sound, or the acoustic feature (Dai *et al*., 2018; Pérez *et al*., 2017). To our knowledge, there have been no studies that directly compare the semantic features and the inter-brain features in a cocktail party scenario. However, the meaning, or the semantic feature is vital to understand the competing speech streams. Recent studies also revealed that processing of semantic or linguistic features could also be modulated by attention, for the features in the attended stream could also better understood than the unattended stream (Broderick *et al*., 2018; Connolly *et al*., 1990; Dai *et al*., 2022; Har-shai Yahav & Zion Golumbic, 2021; Heil *et al*., 2004). It still doesn’t know whether the speaker and listener aligned to each other sole rely on the meaning in the speech or have the extra-linguistic attention modulation (Hartley & Poeppel, 2020; Hasson *et al*., 2012; Pérez & Davis, 2023; Schoot *et al*., 2016; Stolk *et al*., 2016).

The objective of this study was to compare the attention to the speech and the speaker using a cocktail party paradigm. More specifically, we investigated the speaker- listener coupling in contrast to the coupling between the listener’s neural activity and two types of features: the acoustic and semantic features. Naturalistic speech was used as stimulus material, which contains much richer information than either the sound sequence or single word stimulation employed in previous studies. The sound, the meaning, and the speaker could appear at the same time in an ecological situation and enables us to directly compare couplings to the different features simultaneously (Broderick *et al*., 2018; Hamilton & Huth, 2018; Hartley & Poeppel, 2020; Nastase *et al*., 2021; Sonkusare *et al*., 2019; Willems *et al*., 2020). A sequential dual-brain approach(Redcay & Schilbach, 2019) was used, and the electroencephalogram (EEG) of both speaker and listener was recorded. A temporal response function (TRF) method was used to measure the difference between the coupling to attended and unattended features. We also used the neural language processing (NLP) models to help us extract the meaning in the text as the semantic feature. Based on previous studies, we hypothesized that speaker-listener coupling was distinct from the speech-listener coupling, which reveals beyond the speech attention modulation: for the time course, The listener’s neural activity is aligned with the speaker’s neural activity in a broader time range, may have a leading coupling (Dai *et al*., 2018; Kuhlen *et al*., 2012; Stephens *et al*., 2010; Zheng *et al*., 2018). The attention modulation for the acoustic and semantic feature occurs after the speech onset (Ding & Simon, 2012a, 2012b; Hillyard *et al*., 1973; O’Sullivan *et al*., 2015; Power *et al*., 2011; Teoh *et al*., 2022), and the attention effect of the semantic feature lasts longer (Broderick *et al*., 2018; Dai *et al*., 2022). For the frequency bands, we hypothesize that the delta band reflects attention modulation for the inter-brain feature and the semantic feature for they related to the comprehension (Bai *et al*., 2022; Dai *et al*., 2022; Lu, Jin, Ding, *et al*., 2022; Teoh *et al*., 2022; Yu *et al*., 2022; Zuk *et al*., 2021), and the theta band represents the acoustic feature (Ding *et al*., 2014; Etard & Reichenbach, 2019).

## Materials and Methods

### Ethics statement

The study was conducted in accordance with the Declaration of Helsinki and was approved by the local Ethics Committee of Tsinghua University. Written informed consent was obtained from all participants.

### Participants

Two participants (both male, aged 26 and 24 years) were recruited for this study as speakers. Both speakers were from the broadcasting station of Tsinghua University and had experience related to broadcasting and hosting.

Twenty college students (10 females; mean age: 24.7 years; range: 20–43 years) from Tsinghua University participated in the study as paid volunteers for listeners. All participants were native Chinese speakers and reported having normal hearing and normal or corrected-to-normal vision.

The sample size (N = 20) was decided empirically following previous TRF-based studies on human speech processing (Broderick *et al*., 2018; Di Liberto *et al*., 2015; J. Li *et al*., 2022; Mirkovic *et al*., 2015).

### Experimental procedure for the speakers

A sequential inter-brain approach was adopted by the present study (Redcay & Schilbach, 2019), in which the neural activities of the speakers were recorded prior to the listeners. The sequential design was more appropriate for this study than the real-time interactive design because the speakers’ audio and neural activity remained consistent for all listeners (Leong *et al*., 2017; Y. Liu *et al*., 2017; Stephens *et al*., 2010).

In this experiment, each speaker participated in 30 trials, each of which was approximately 51–76 seconds in length, while the speakers’ audio signals and EEG signals were recorded. The experimenter selected 28 trials for the listener’s experiment, excluding the two most unqualified trials.

The speaker first read the relevant material on the screen. There was a wide variety of content to be covered, including one’s hometown, a recent book, a fable, etc. The speaker could decide how long they wanted to spend on preparation and start talking when they were fully prepared (the length of preparation was usually 3 minutes). When the speaker was prepared, they would press the spacebar on the computer keyboard, and the recording would begin. When the spacebar was pressed, three 1000 Hz pure tone cues were triggered (duration: 1000 ms; cue interval: 1500 ms). The cues were presented as event markers, synchronized with the sound in the listener’s experiment to ensure that the neural signals of the speaker and listener remain aligned with the sound stimuli. The speaker was asked to start speaking immediately after the end of the third beep (within approximately 3 s). A fixation and a countdown timer appeared on the screen during the talking part. The speaker was asked to stare at the fixation and to complete the speaking as clearly, completely, and naturally as possible. During the recording process, the experimenter listened to the speaker’s narration simultaneously and controlled the quality. The experimenter had the right to ask the speaker to retell the clip if there was a reason for the lack of fluency, length, etc., that might affect the listeners’ perception. The materials of both speakers’ content were varied. Between each trial, the speakers were allowed to rest on their own. During the experiment, the speakers were asked to control their head movements and facial muscles to obtain better quality EEG signals.

The speech stimuli were recorded from two male speakers using the microphone of an iPad2 mini (Apple Inc., Cupertino, CA) at a sampling rate of 44,100 Hz.

### Experimental procedure for the listeners

The experiment consisted of four blocks, each containing seven trials. Two speech streams were presented simultaneously in each trial, one to the left ear and the other to the right ear. Two speech streams of the same trial matched the volume, i.e., the root mean squared intensity of the amplitude of the speech streams in the same trial were the same. The participants were instructed to attend to one spatial side according to the hints on the screen (“Please pay attention to the [LEFT/RIGHT]”). Considering the possible duration difference between the two audio streams, the researchers set the end of the trial after the longer speech audio had ended. Each trial began when participants pressed the SPACE key on the computer keyboard. A white fixation cross was also displayed throughout the trial. The speech stimuli were played immediately after the keypress and were preceded by the three beep sounds.

At the end of each trial, four multiple-choice questions (two for the attended story and the other two for the unattended story) were presented sequentially in random order on the computer screen. Each question had four options, and participants entered the letter of the correct option as their answer. The listeners were not explicitly informed about the correspondence between the questions and the stories. For instance, one question following a story about one’s hometown was, “What is the most dissatisfying thing about the speaker’s hometown? (推测讲述人对于家乡最不满意的地方在于?)”, and the four choices were A) There is no heating in winter; B) There are no hot springs in summer; C) There is no fruit in autumn; D) There are no flowers in spring (A. 冬天 没暖气; B. 夏天没温泉; C. 秋天没水果; D. 春天没鲜花). The single-trial comprehension accuracy could be 0% (two wrong answers), 50% (one correct answer), or 100% (two correct answers) for both the attended and the unattended stories. No feedback was given as to whether the questions were answered correctly.

After completing these questions, participants rated their concentration level of the attended stream, the experienced difficulty performing the attention task, and the familiarity with the attended material using three 10-point Likert scales. Throughout the trial, participants were required to maintain visual fixation on the fixation cross while listening to the speech. Meanwhile, they were asked to minimize eye blinks and all other motor activities. The participants were recommended to take a short break (around 1 min) after every trial within one block and a long break (no longer than 10 min) between blocks.

In each block, the side being attended to was fixed (two blocks for attending to the left side and two for attending to the right side). Within each block, the identity of the speaker is kept constant on the left and right sides. The to-be-attended spatial side and the corresponding speaker identity were balanced within the participant, with seven trials per side for both speakers. The assignment of the stories to the four blocks was randomized across the participants.

The experiment was conducted in a sound-attenuated, dimly lit, and electrically shielded room. The participants were seated in a comfortable chair in front of a 19.7- inch LCD monitor (Lenovo LT2013s). The viewing distance was approximately 60 cm. The experimental procedure was programmed in MATLAB using the Psychophysics Toolbox 3.0 extensions (Brainard & Brainard, 1997). The speech stimuli were delivered binaurally via an air-tube earphone (Etymotic ER2, Etymotic Research, Elk Grove Village, IL, USA) to avoid possible electromagnetic interference from auditory devices. The volume of the audio stimuli was adjusted to be at a comfortable level (∼70 dB SPL) that was well above the auditory threshold. The average presentation level was measured with a B&K (Brüel & Kjær, Nærum, Denmark) Sound Level Meter (Type 2250 Investigator) with a 1-inch Free-field Microphone (Type 4144) and an Artificial Ear (Type 4152).

### Data acquisition and pre-processing

EEG was recorded from 60 electrodes (FP1/2, FPZ, AF3/4, F7/8, F5/6, F3/4, F1/2, FZ, FT7/8, FC5/6, FC3/4, FC1/2, FCZ, T7/8, C5/6, C3/4, C1/2, CZ, TP7/8, CP5/6, CP3/4, CP1/2, CPZ, P7/8, P5/6, P3/4, P1/2, PZ, PO7/8, PO5/6, PO3/4, POZ, Oz, and O1/2), which were referenced to an electrode between Cz and CPz, with a forehead ground at Fz. A NeuroScan amplifier (SynAmp II, NeuroScan, Compumedics, USA) was used to record EEG at a sampling rate of 1000 Hz. Electrode impedances were kept below ten kOhm for all electrodes.

The recorded EEG data were subjected to an artifact rejection procedure using independent component analysis. Independent components (ICs) with large weights over the frontal or temporal areas, together with a corresponding temporal course showing eye movement or muscle movement activities, were removed. The remaining ICs were then back-projected onto the scalp EEG channels, reconstructing the artifact-free EEG signals. While the relatively long duration of the speech trials in the present study (∼1 minute per story, see Experimental procedure) has made it more difficult for the participants to avoid inducing movement-related artifacts as compared to the classical ERP-based studies, a temporally continuous, non-interrupted EEG segment per trial was preferred for the employment of the ICA method. Therefore, any ICs with artifact-like EEG activities for more than 20% of the trial time (i.e., ∼12 sec) were rejected, leading to around 4–11 ICs rejected per participant. The cleaned EEG data were used for the mTRF analysis without any further artifact rejection procedures.

Next, the EEG data were segmented into 28 trials according to the markers representing speech onsets. The analysis window for each trial was extended from 10 to 55 s (duration: 45 s) to avoid the onset and the offset of the stories.

The pre-processed EEG signals were re-referenced to the average of all scalp channels and then downsampled to 128 Hz before the modeling. Then, the EEG data were filtered in delta (1–4 Hz) and theta (4–8 Hz) (filter order: 64, one-pass forward filter). The use of a causal FIR filter ensured that filtered EEG signals were decided only by the current and previous data samples (de Cheveigné & Nelken, 2019), which is essential for accurate time-course analysis. The filter order of 64 was chosen to keep a balance of temporal resolution and filter performance: the filtered EEG signals were therefore calculated based on the preceding 500 ms data (64 at 128 Hz).

### Speaker-listener coupling: Inter-brain Features

The inter-brain method enables us to study how the listeners are aligned with the attended speaker (Dai *et al*., 2018; Hasson *et al*., 2012; Jing Jiang *et al*., 2021; Stephens *et al*., 2010). We used the speaker’s neural activity as the representation of the inter-brain feature. The speaker’s EEG served as the inter-brain feature. It followed the same pre-processing procedure as the listener’s EEG.

### Speech listener coupling: acoustic and semantic features

- *Acoustilistc Features*

The amplitude envelope of the speech represented the acoustic features of the speech. It was obtained using a Hilbert transform and then down-sampled to the same sampling rate of 128 Hz.

- *Semantic Features*

The original audio recorded by the speaker during the EEG recording was converted to the text first automatedly by *Iflyrec* software (Iflytek Co., Ltd, Hefei, Anhui) and then double checked manually. The onset time of every word was extracted during this process.

The recent emergence of Natural Process Language (NLP) models has enabled the description of the semantic features in speech (Brodbeck *et al*., 2022; Broderick *et al*., 2018, 2021). Next word prediction is one of the fundamental NLP tasks using the semantic information in the texts (Schrimpf *et al*., 2021; Vaswani *et al*., 2017). The goal of the task is to predict the next word when given a sequence of words *W*_1_, *W*_2_, …, *W*_t–1_, which was consistent with the human understanding process. The probability of the next word is *P*(*W_t_*|*W*_1_, …, *W*_t–1_) and can be calculated by varied NLP models. The surprisal of the word was defined as follow, which reflected how surprised the next word (Willems *et al*., 2016):

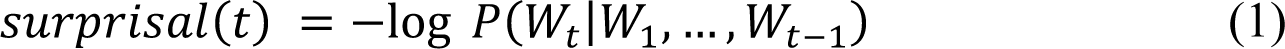

The index was calculated based on ADAM, a widely accepted classical natural language process model (Bengio *et al*., 2003; Kingma & Ba, 2015). The model was trained on a couple of *People’s Daily.* There were 534,246 words involved in the model training. 66,781 words were in the cross-validation set, and 66,781 words used as a test set. The details of the model are described in Table 1.

**Table 1.** Details of Neural Language Processing Model

| Type | Parameter |
| --- | --- |
| model type | LSTM |
| embedding size | 200 |
| hidden units per layer | 200 |
| number of layers | 2 |
| initial learning rate | 3 |
| gradient clipping | 0.25 |
| sequence length | 35 |
| drop out | 0.2 |
| epoch | 50 |
| batch size | 3 |

After calculating the surprisal index of every word, we generated a “semantic vector” at the same sampling rate as the EEG data (Broderick *et al*., 2018). The vectors contained the time-aligned impulses at the start of each word of the surprisal value for every audio clip.

### Temporal response function modeling

The analysis workflow for the analysis related to the attended speech stream is shown in Figure 1. The neural responses to the three different features were characterized using a temporal response function (TRF)-based modeling method (Crosse *et al*., 2016, 2021).

**Figure 1.**
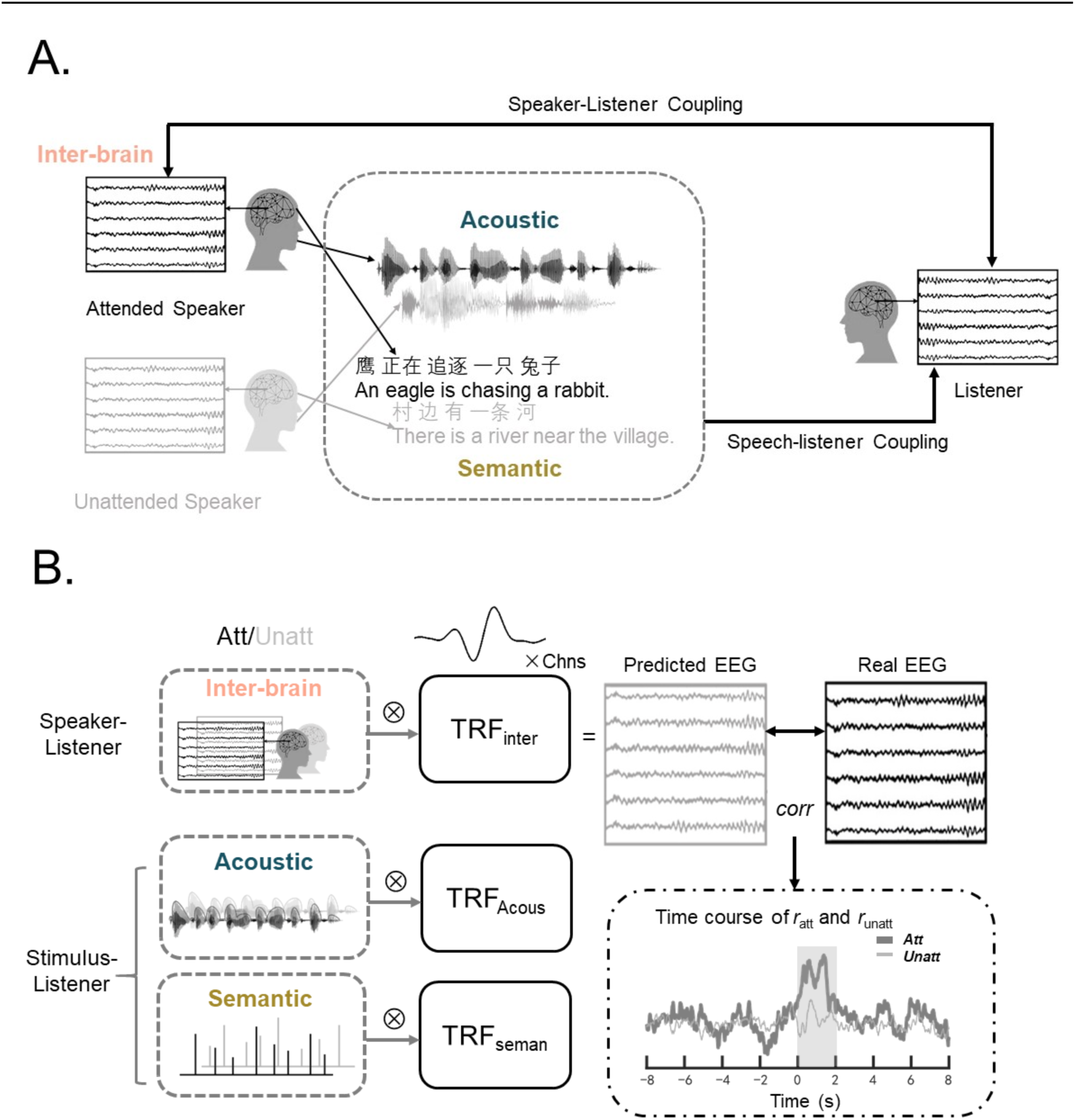
The stimuli, the experimental paradigm and the analysis process. (A) The paradigm and the different features. A “cocktail party” selective attention paradigm was used, in which the listener was asked to pay attention to one side of the speech stream and ignore the other. While listening to the speech stream, the listener’s EEG signals were recorded. The speaker’s EEG signals were defined as the “inter-brain” features in our study. The speaker-listener coupling is “beyond the stimulus”. The acoustic and semantic features were included in the coupling between the speech stimuli and the listener. (B) The data analysis process. The encoding models were trained for different features separately. The Encoding *r* for the attended feature and the unattended feature was calculated time point by time point by applying the TRF method. The TRF of each feature was estimated separately in two frequency bands: delta and theta.

Three different features mentioned above are the input signal required by TRF. The corresponding neural response *r*(*t, n*) can be formulated as follows:

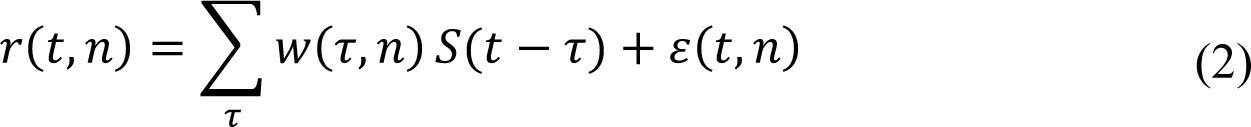

where *r*(*t, n*) is the actual EEG response at every channel *n* ; *t* = 1 … *T* is the time point; *S*(*t* − *τ*) means the multivariate stimulus representation; *W*(*τ*, *n*) is the channel specific TRF at lag *τ* and *ε*(*t*, *n*) is the residual.

The TRF is estimated by minimizing the mean square error between the actual neural response *r*(*t*, *n*) and the neural response predicted by the model *ř*(*t*, *n*). The Pearson’s correlation between the actual neural response and predicted neural response was referred as Encoding *r.* The mTRF toolbox (Crosse *et al*., 2016) was used to estimate the TRF(*w*) as fellow:

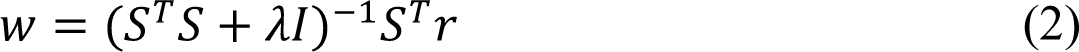

where λ is the ridge regression parameter, *I* is the identity matrix, and the matrix *S* is the stimulus matrix. The lambda varied from 10^-1^ to 10^8^ (lambda = 10^-1^, 10^0^, …, 10^8^) to make the model optimal (Crosse *et al*., 2021; Teoh *et al*., 2022). The λ value, which produces the highest encoding *r*, averaged across trials and channels, was selected as the regularization parameter for all trials per participant (Broderick *et al*., 2019). The cross-validation procedure was implemented in a leave-one-trial-out manner: the TRFs were trained based on data from 27 trials and tested on the left-out trial each time.

The TRF was trained at every individual time lags of -8 s to 8 s to investigate the specific interval of attentional modulation for each feature. At a sampling rate of 128 Hz, there are 2049 individual time-lag intervals of 7.625 ms. The choice of the window was aligned with the previous fMRI and fNIRS inter-brain studies.

The TRF calculation procedure was performed for the EEG signals from each EEG channel filtered at the delta and theta frequency bands. Only attended features are used as input to the model, and TRFs trained by the attended features and the neural response were applied to the tests of attended and unattended features, referred to as Encoding Encoding *r*_att_ and Encoding *r*_unatt_, respectively. This approach was largely used in the attention-related studies (Mirkovic *et al*., 2015; O’Sullivan *et al*., 2015).

### Quantification and statistical analysis

The paired *t*-tests were performed to investigate the attentional modulation for different features, contrasting the encoding *r* of the attended speech versus the unattended speech at each time lag. The Fisher-*z* transformation was used to convert *r* encoding into *z*-score (Corey *et al*., 1998). The transformation transformed the *r* values into a normal distribution that can be compared and assessed for statistical significance.

A nonparametric cluster-based permutation analysis was applied to account for multiple comparisons (Maris & Oostenveld, 2007). In this procedure, neighboring channel-latency bins with uncorrected *t*-tests *p*-value below 0.05 were combined into clusters, for which the sum of the correlational *t*-statistics corresponding to the *t*-tests was obtained. The combining process was initially automated by the toolbox and then manually double-checked. Two clusters were combined if they shared a similar spatial distribution (e.g., the clusters have a large overlap of the electrons). A null distribution was created through permutations of data across participants (*n* = 1,000 permutations), which defined the maximum cluster-level test statistics and corrected *p*-values for each cluster. Clusters with *p*-values below 0.01 based on clusters were selected for further analysis.

The above statistical analysis followed the standard cluster-based permutation procedure as employed in classical ERP and related studies (Arnal *et al*., 2011; Henry & Obleser, 2012; Zhang *et al*., 2012). Note that the reported *p*-values were only corrected for the tests performed within each frequency band by using cluster-based permutation tests. No multiple comparison correction was employed across different frequency bands.

### Correlation between clusters

The *Spearman* correlation of Encoding *r*_att_ in each cluster was calculated for each pair of clusters to analyze the correlation between them.

### Partial correlation between the behavioral performance and the neural activity

The partial correlation between the behavioral performance and the coefficients in was calculated to reveal the unique contribution of a certain feature to the comprehension performance. Specifically, the correlations between the mean Encoding *r*_att_ in the cluster and the accuracy of the questions were calculated while controlling other features.

### Data Availability Statement

Data generated in our study has been uploaded to Mendeley Data (Li, Jiawei (2022), “Speaker-listener Cocktail Party”, Mendeley Data, V3, doi: 10.17632/wkh75s27kn.3).

When existing toolboxes were used for data analysis, citations have been provided.

## Results

### Behavioral Performance of the Listeners

The average comprehension performance was significantly better for the 28 attended stories than for the 28 unattended stories (67.0±2.5% (standard error) vs. 36.0±1.6%, *t*(19) = 10.948, *p* < .001; the four-choice chance level: 25%). The participants reported a moderate level of attention (8.146±0.343 on a 10-point Likert scale) and attention difficulties (2.039±0.530 on a 10-point Likert scale). The accuracy for the attended story was significantly correlated with both the self-reported attention level (*r* = .476, *p* = .043) and attention difficulty (*r* = -.677, *p* = .001). The self-reported story familiarity level was low for all the participants (0.860±0.220 on a 10-point Likert scale) and was not correlated with comprehension performance (*r* = -.224, *p* = .342). These results suggest that participants’ selective attention was effectively manipulated, and the measurement of comprehension performance was reliable. The response accuracy varied from 25.0 % to 51.8% for unattended stories.

### The speaker-listener coupling precedes the onset of speech on delta band

A cluster-based permutation (Maris & Oostenveld, 2007) was conducted to reveal the difference between the encoding of the attended and the unattended features and control for multiple comparisons. As figure 2 shows, there was only one inter-brain cluster found in the delta band (cluster-based *p* < .001). The inter-brain cluster had a wide time range of -4.836 to -0.539 s with the electrodes in the left frontal region. Figure S1 displays results on other frequency bands.

**Figure 2.**
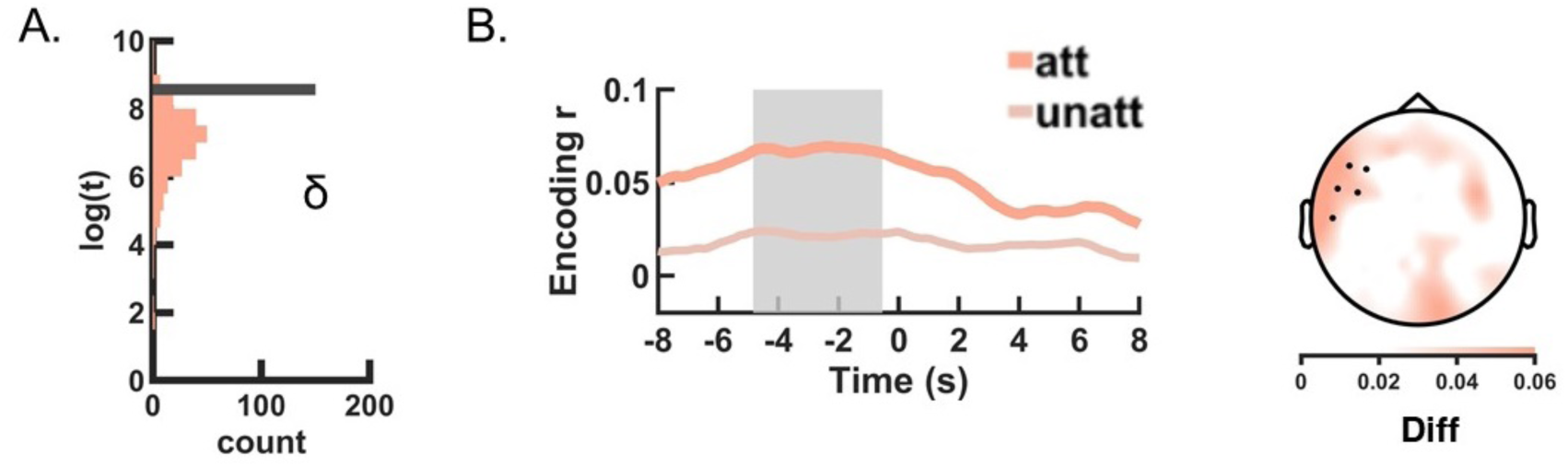
The attention modulation in speaker-listener coupling. (A) The null distribution of the cluster-based log(*t*-statistics) of the inter-brain feature. For illustration purposes, the *t*-statistics were transformed into log(*t*). The label “count” for the *x*-axis shows the number of occurrences of the corresponding *t*-value in the 1000 permutations. The gray lines indicate significant clusters. (B) The time course and the topo-plot of the significant inter-brain cluster. The light pink line represents the Encoding *r_att_*, and the dark pink represents the Encoding *r_unatt_*. The shaded region depicts a significant difference in the time window. The topo-plot of the average difference between Encoding *r_att_* and Encoding *r_unatt_* cluster. The black dots indicate the channels in the cluster.

### The attention modulation for acoustic and semantic features occurs after the speech onset

As shown in Figure 3, the *Acoustic_Early* cluster covered the left-lateralized fronto-central and occipital electrodes (cluster-based permutation *p* < .001) at a latency of 0.219−0.359 s after the onset of speech. The *Acoustic_Late* cluster had a later latency of 1.508−1.602 s with the electrodes in the right-frontal regions (cluster-based *p* = .005). The clusters only appeared on the theta bands.

**Figure 3.**
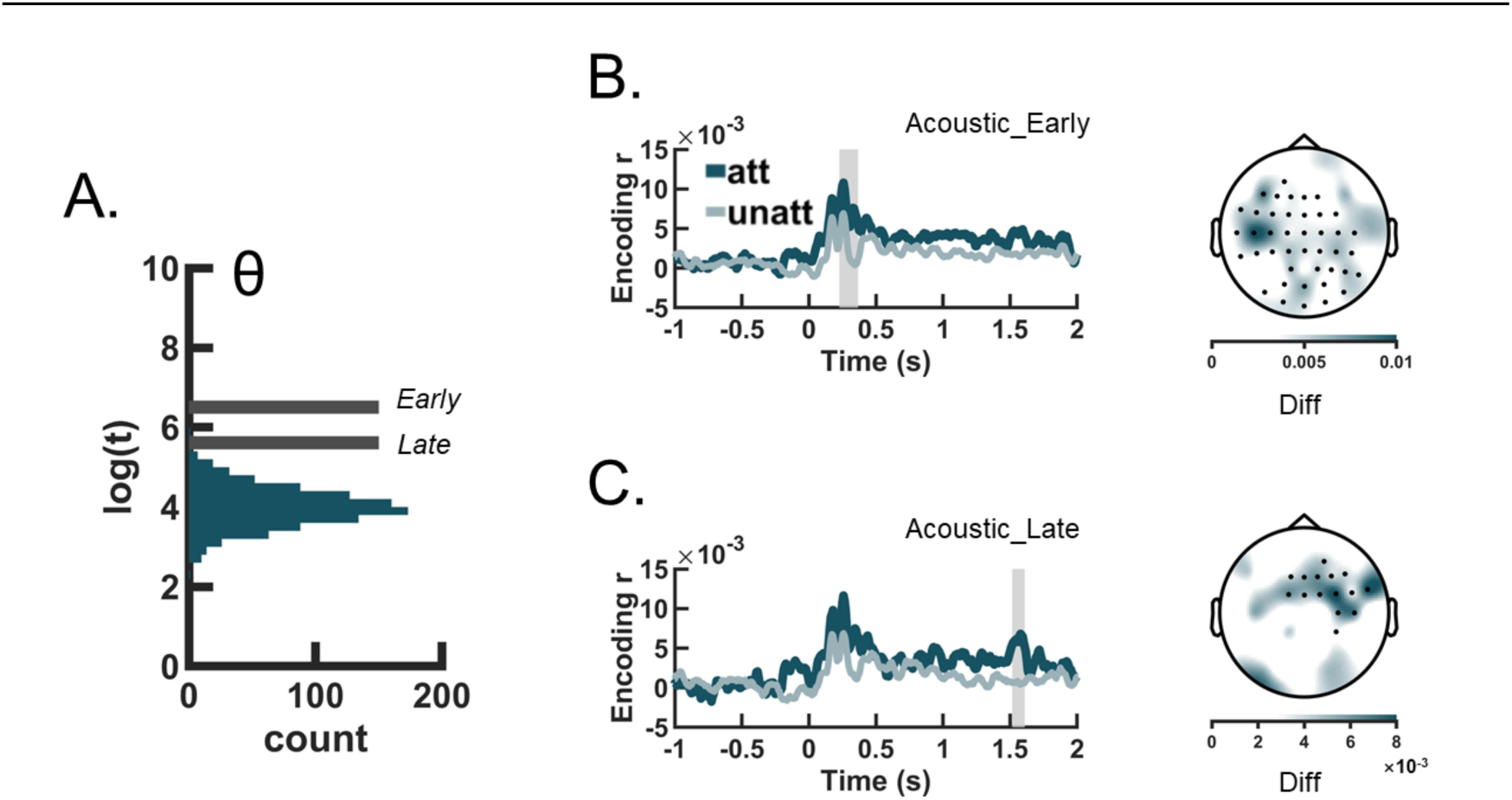
The attention modulation for the acoustic feature. (A) The null distribution of the cluster-based log(*t*-statistics) of the acoustic feature. (B) and (C) The time course and the topo-plot of the significant acoustic cluster. The dark blue line represents the Encoding *r_att_*, and the light line represents the Encoding r*_unatt_*. The shaded region depicts a significant difference in the time window. The topo-plot of the average difference between Encoding *r_att_* and Encoding *r_unatt_* cluster. The black dots indicate the channels in the cluster.

In contrast to the acoustic feature, semantic clusters were found in the delta band. As Figure 4 indicates, the *Semantic_Early* cluster occurred at 0.227−0.621 s covering the electrodes in frontal and central regions (cluster-based *p* = .002). The *Semantic_Late* cluster was found at 1.073−1.516 s involving the wide distribution of the electrodes (cluster-based *p* < .001).

**Figure 4.**
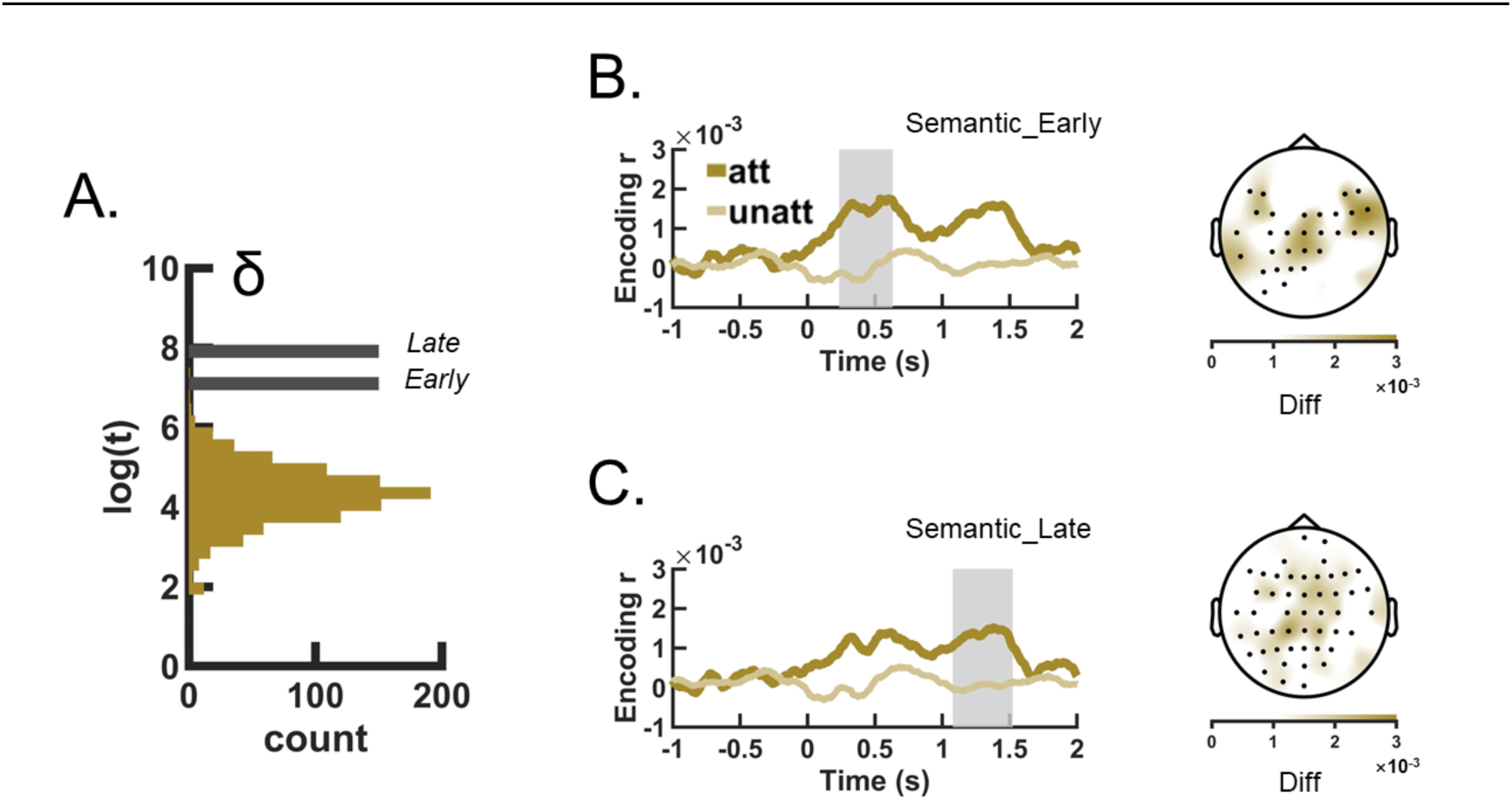
The attention modulation for the semantic feature. (A) The null distribution of the cluster-based log(*t*-statistics) of the semantic feature. (B) and (C) The time course and the topo-plot of the significant semantic cluster. The dark brown line represents the Encoding *r_att_*, and the light brown represents the Encoding *r_unatt_*. The shaded region depicts a significant difference in the time window. The topo-plot of the average difference between Encoding *r_att_* and Encoding *r_unatt_* cluster. The black dots indicate the channels in the cluster.

### The speaker-listener coupling was correlated to the comprehension performance

As Table 2 illustrated, only the inter-brain cluster has a significant partial correlation with the behavioral performance, partial correlation *r* = -.769, *p* = .002 (FDR corrected, Figure 5B). All the other clusters didn’t reveal significant correlations. We further analyzed the Encoding *r*_att_ in the inter-brain cluster and difficulty. We found significant positive correlation, *r*= .499, *p* = .025 as shown in Figure 5C.

**Figure 5.**
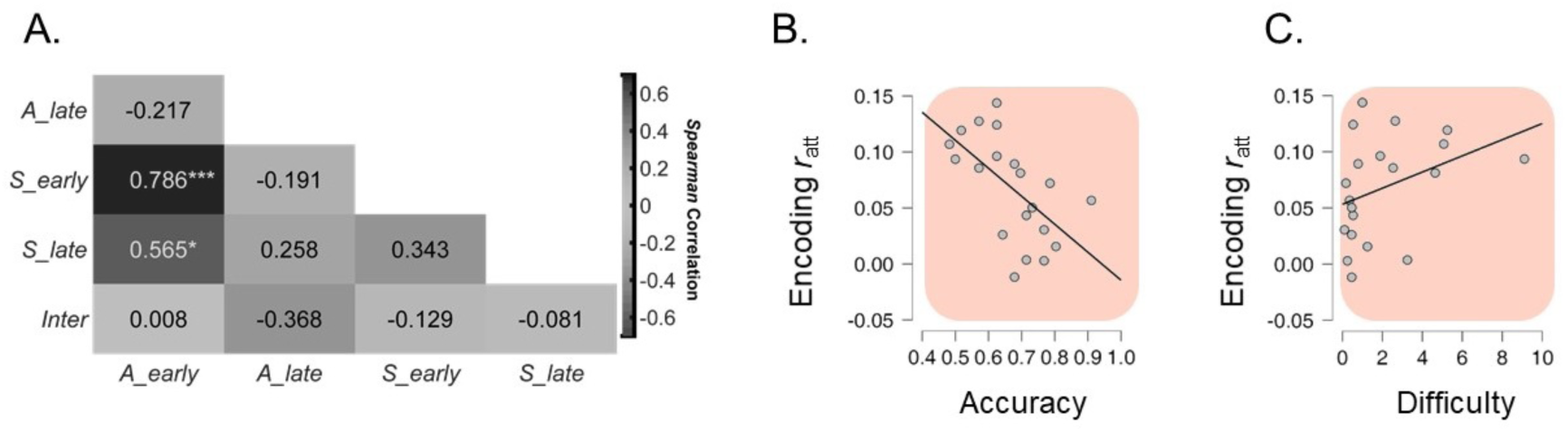
The correlation between the neural activity and the behavioral result. **(A)**The pairwise *spearman* correlations between Encoding *r*_att_ in different clusters. The acoustic features and the semantic features correlated with each other. *A_early* stands for Acoustic_Early, *A_late* stands for Acoustic_Late, *S_early* is short for Semantic_early, *S_late* stands for Semantic_Late, and *Inter* is short for Inter-brain. **(B)** and **(C)** The correlation between the Encoding *r*_att_ in inter-brain clusters and the accuracy of the comprehension questions and the self-rated difficulty.

**Table 2.** The partial correlation between the Encoding r and accuracy of the comprehension performance

| Clusters | $\rho$ | $p$ values (FDR-corrected) |
| --- | --- | --- |
| Acoustic-early | .512 | .089 |
| Acoustic-late | -.306 | .291 |
| Semantic-early | -.412 | .170 |
| Semantic-late | .098 | .710 |
| Inter-brain | -.769 ** | .002 |

\* $p < .05$ , \*\* $p < .01$ (FDR corrected)

As Figure 5A indicates, the average Encoding *r*_att_ in *Semantic_Early* cluster and the average Encoding *r*_att_ in *Acoustic_Early* clusters were highly correlated (*r*(18) = .786, *p* < .001, FDR_corrected). The average Encoding *r*_att_ in *Semantic_late* cluster and the average Encoding *r*_att_ in *Acoustic_Early* cluster were also highly correlated (*r*(18) = .565, *p* = .045, FDR_corrected). There were no other significant correlations between other clusters (*p*s > .05).

## Discussion

Our study directly compares the speaker-listener coupling and the stimulus-listener coupling in the “cocktail party scene” using the high temporal resolution EEG inter-brain recording method. Our result uncovered a distinctive pattern of neural alignment, wherein the listener synchronized with the speaker and speech stimulus in disparate ways: The listeners aligned to the speaker’s neural activity 5 s before the speech onset, which reveals a significant difference from the speech-listener coupling. Other two types of couplings occur after the speech onset: The attention modulated the sound 200−350 ms after the speech onset in the theta band. The meaning of speech was modulated later, in the delta band at 200−600 ms.

Our study proved the distinct role of the attentional speaker-listener coupling in the “cocktail party”: the spatial distribution and the time range are different from the couplings to the acoustic feature and the semantic feature. For the spatial distribution, the central electrodes are primarily involved in the acoustic and semantic modulations, only the left-frontal electrodes are recruited in the inter-brain modulation. The left frontal regions play a critical region in the language process (Har-shai Yahav & Zion Golumbic, 2021; Hickok & Poeppel, 2007) and are a ‘high-order’ area in attention selection (Zion Golumbic *et al*., 2013). The left-frontal electrodes may indicate a unique contribution of IFG when listeners are under adverse listening conditions in previous inter-brain studies (Dai *et al*., 2018; Z. Li *et al*., 2021; L. Liu *et al*., 2020). For the time range, the listeners are aligned to the speaker’s neural activity 5 s before the speech onset, where other two features occur after the speech. This result replicated previous inter-brain studies using fMRI or fNIRS in the high temporal resolution EEG signals recording method (Dai *et al*., 2018; J. Jiang *et al*., 2012; L. Liu *et al*., 2020; Stephens *et al*., 2010). Our result further suggests that the speaker-listener coupling was different from speech-listener coupling, which highlights the significance of the speaker in the speech comprehension process (Dai *et al*., 2018; Pérez *et al*., 2017, 2019).

The correlation to the behavioral performance further highlights the unique role of the speaker-listener coupling. In our study, the coupling to the acoustic feature and semantic feature were correlated, but the coupling to the speaker’s neural activity is independent of the speech stimuli. The speaker-listener coupling had a negative correlation with the comprehension performance (Figure 5B). We further found that the neural response was positively correlated with the perceived task difficulty reported by the listeners (Figure 5C). We successfully replicated the findings of previous inter-brain studies that indicate a relationship between inter-brain coupling and comprehension performance (Z. Li *et al*., 2021, 2022; Stephens *et al*., 2010). However, in contrast to these studies, our results reveal a negative correlation. The discrepancy observed may be attributed to variations in the recording methodology employed: the previous studies using mainly the blood oxygen signals, like fNRIS and fMRI, but we used a high temporal resolution EEG signal. The coupling in two types of signals may have different interpretations. We called the negative correlation observed in our study a “compensation” mechanism: when the listeners find it hard to complete the task, they start to guess what the speaker may want to say. However, guessing is not always correct. Therefore, the comprehension performance is decreasing. While speech itself may only serve as a trigger and an entrainment signal, the listener could actively “understand” the speaker not only through the speech itself but rely on the “beyond the stimulus” grounding (Bashivan *et al*., 2019; Hartley & Poeppel, 2020; Hasson *et al*., 2012; Jing Jiang *et al*., 2021; Redcay & Schilbach, 2019; Stolk *et al*., 2016).

Our study reveals the distinct roles of delta and theta bands in attentional modulation. The theta band modulates with the acoustic feature, and the delta band modulates with the semantic feature and inter-brain feature. This result possibly extends our understanding of the functional roles of the two bands. It is consistent with the previous finding that the theta band processes stimulus-linked features, like the acoustic feature (Ding *et al*., 2014; Ding & Simon, 2012a, 2012b; Teng *et al*., 2017), and the delta band reflects comprehension-related features, like the semantic feature (Ding *et al*., 2014; Etard & Reichenbach, 2019; J. Li *et al*., 2022; Lu, Jin, Pan, *et al*., 2022). In particular, the modulation for the inter-brain feature was also found in the delta band, which may indicate that the delta bands reflect the comprehension-related functions.

Our study introduces novel analytical methods to the field of inter-brain studies. We applied the temporal response function method (TRF) to depict the listener’s neural response to a high dimensional speaker’s EEG signal. This method was widely used in the studies of naturalistic speech (Crosse *et al*., 2016; Di Liberto *et al*., 2015; Ding & Simon, 2012a, 2012b; O’Sullivan *et al*., 2015; Power *et al*., 2012), as well as the visual studies (Jia *et al*., 2017, 2019) and memory studies (Huang *et al*., 2018). Using the TRF method enables a direct comparison between the speaker-listener coupling and different types of speech-listener coupling. We also used the NLP models (Armeni *et al*., 2019; Brodbeck *et al*., 2022; Broderick *et al*., 2018; Kingma & Ba, 2015) to describe the semantic meaning in the speech as the latest review suggested (Jing Jiang *et al*., 2021; Pérez & Davis, 2023). The combined use of different lately methods enhance our understanding of the attention modulation in a comprehensive way and provide a new inspiration for the future studies underlying inter-brain coupling.

While this study sheds light on a new perspective in the “cocktail party effect”, there are several areas that warrant further exploration. Firstly, our study simplified the semantic feature into the “surprisal index”, which is a common choice in previous studies(Aurnhammer & Frank, 2019; Brodbeck *et al*., 2022; Heilbron *et al*., 2022; Mesik *et al*., 2021; Willems *et al*., 2016). However, there were alternative approaches to extract the semantic feature in the text, such as the word embeddings. Subsequent studies could undertake a comparative analysis of various semantic representations, such as surprisal, entropy, or word embeddings, generated by diverse models including word2vec (Broderick *et al*., 2018; Lu, Jin, Pan, *et al*., 2022; Mikolov *et al*., 2013), GPT (Heilbron *et al*., 2022; Schramowski *et al*., 2022; Solaiman *et al*., 2019) or BERT (Devlin *et al*., 2019). This would enable a comprehensive evaluation of the consistency of their findings. Meanwhile, our studies used a sequential-dual-brain approach to use the speaker-listener coupling. While this approach has the potential to reduce variance originating from the speaker’s neural activity (Leong *et al*., 2017; Z. Li *et al*., 2021; Redcay & Schilbach, 2019), it may not fully capture the complexity of neural activity present in real-life conversations (Dai *et al*., 2018; Jing Jiang *et al*., 2012; Pérez *et al*., 2017). Further studies could also apply the simultaneous dual-brain approach (Redcay & Schilbach, 2019) to further investigate the issue.

In conclusion, our study used the attention as a spotlight and revealed that the listener would strike the same “neural notes” with the speaker before the speech onset, while the acoustic note is struck on the theta bands after the speech onset firstly, and the semantic note comes later and lasts longer on the delta band. Our study depicts the temporal dynamics of the attention modulation and the functional roles of different frequency bands, which contributes to the old “cocktail party” a new integrative perspective. In an era of rising artificial intelligence, our study still highlights the value and significance of human-to-human interaction between speakers and listeners, as the result demonstrates the possibility of mutual understanding between the speaker and listener “beyond the stimulus” in their neural activities (Yeshurun *et al*., 2021).

## Acknowledgments

This work was supported by the National Science Foundation of China (NSFC) and the German Research Foundation (DFG) in project Crossmodal Learning (grant number: NSFC 62061136001/DFG TRR-169/C1, B1), the National Natural Science Foundation of China (grant number: 61977041) and a postdoctoral fellowship of the Humboldt Foundation to J.L.

The authors would like to thank Prof. Dr. Xiaoqin Wang and Dr. Yue Ding for providing the shielded room for the experiment as well as necessary technical support.

The authors would like to express their gratitude towards Dr. Zhiyuan Liu and members of his lab for computing the NLP models.

## Notes

### Competing Interest Statement

The authors have declared no competing interest.

### Summary of Updates

The manuscript has been updated with a substantial revision on the Introduction part, to focus on the inter-brain results.

