## Supplemental Figure S1 for "EEG-based speaker-listener neural coupling reflects speech-selective attentional mechanisms beyond the speech stimulus"

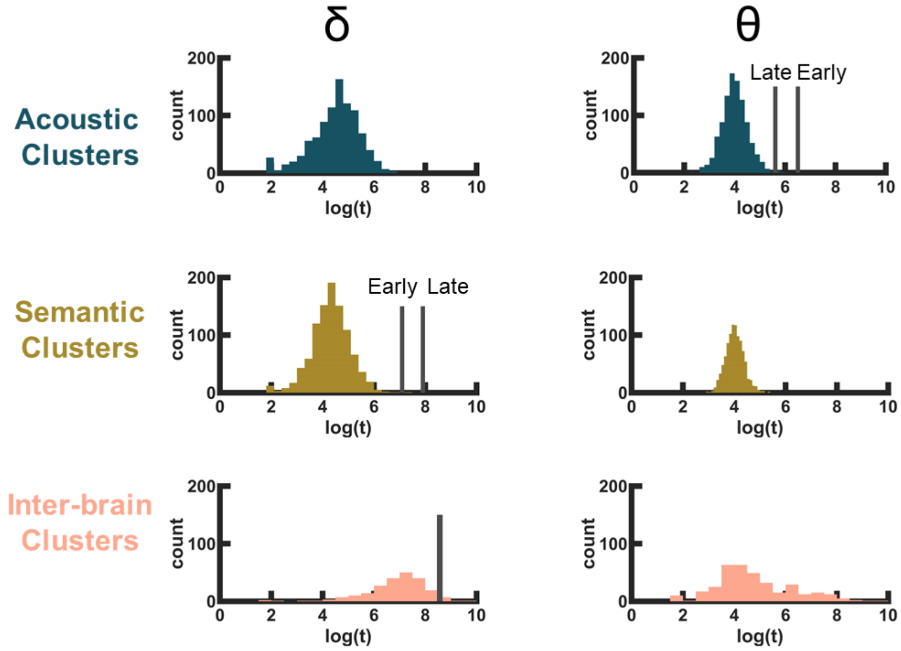


**Figure S1 Different attentional modulation roles of different bands.** The null distribution of the *t*-statistics of every feature in the delta band(left) and theta band(right). For illustration purposes, the *t*-statistics were transformed into log(*t*). The label "count" for the y-axis shows the number of occurrences of the corresponding *t*-value in the 1000 permutations. The grey lines indicate significant clusters.
